# VGLUT3-p.A211V variant fuses stereocilia bundle and elongates synaptic ribbons in the human deafness DFNA25

**DOI:** 10.1101/2020.06.26.170852

**Authors:** Yuvraj Joshi, Stéphanie Miot, Marie Guillet, Gaston Sendin, Jérôme Bourien, Jing Wang, Rémy Pujol, Salah El Mestikawy, Jean-Luc Puel, Régis Nouvian

## Abstract

DFNA25 is an autosomal-dominant and progressive form of human deafness caused by mutations in the *SLC17A8* gene, which encodes the vesicular glutamate transporter type 3 (VGLUT3). To resolve the mechanisms underlying DFNA25, we studied the phenotype of the mouse harboring the p.A221V mutation in human (corresponding to p.A224V in mouse). Using auditory brainstem response and distortion products of otoacoustic emissions, we showed that VGLUT3^A224V/A224V^ mouse replicates the DFNA25 progressive hearing loss with intact cochlear amplification. Scanning electron microscopy examinations demonstrated fused stereocilia bundle of the inner hair cells (IHCs) as the primary cause for DFNA25. In addition, the IHC ribbon synapses undergo structural and functional modifications at later stages. Using super-resolution microscopy, we observed oversized synaptic ribbons associated with an increase in the rate of the sustained releasable pool of exocytosis. These results indicate that the primary defect in DFNA25 stems from a failure in the mechano-transduction followed by a change in synaptic transfer. The VGLUT3^A224V/A224V^ mouse model opens the way to a deeper understanding and to a potential treatment of DFNA25.

## Introduction

Mutations in the *SLC17A8* gene are associated with DFNA25, an autosomal-dominant and progressive form of human deafness (Ruel *et al*, 2008; Ryu *et al*, 2016). *SLC17A8* encodes the vesicular glutamate transporter type 3 (VGLUT3) which belongs to the VGLUTs family responsible for glutamate loading into the synaptic vesicles, a critical step to achieve synaptic transfer (Gras *et al*, 2002; Fremeau *et al*, 2002; Takamori *et al*, 2002). VGLUT3 is expressed in small subsets of neurons in the central nervous system and in the inner ear (Ruel *et al*, 2008; Seal *et al*, 2008; Zhang *et al*, 2011; see for review Mestikawy *et al*, 2011). In zebrafish, VGLUT3 is expressed in the ear and lateral line organ, especially in the hair cells, which convert the mechanical stimuli into glutamate release onto afferent fibers. While the loss of VGLUT3 does not affect the microphonic receptor, reflecting the mechano-transducer current at the hair cell stereociliary bundle, it results in a strong decrease of the posterior lateral line ganglion activity (Obholzer *et al*, 2008). Accordingly, the reduction in the VGLUT3 expression abolishes the vestibulo-ocular and acoustic startle reflexes (Obholzer *et al*, 2008). In mouse, the inner hair cells (IHCs), which transduce the mechano-transduction into glutamate secretion, abundantly express VGLUT3 (Ruel *et al*, 2008; Seal *et al*, 2008), in contrast to the outer hair cells (OHCs), which amplify the acoustic input within the cochlea. The genetic ablation of *SLC17A8* leads to the calcium-triggered exocytosis of empty synaptic vesicles, making the auditory afferent fibers silent (Ruel *et al*, 2008; Seal *et al*, 2008). Thus, the loss of VGLUT3 in mice results into the lack of auditory brainstem responses, reflecting a failure in the ascending auditory pathway activation, and leaves unaffected the distortions products of otoacoustic emission (DPOAEs), as a signature of the OHCs activity (Ruel *et al*, 2008; Seal *et al*, 2008). In addition, it has been reported that the ribbon bodies, the organelles surrounded by a monolayer of synaptic vesicles, are abnormally thin and elongated in the VGLUT3 knock-out (KO) IHC synapses (Seal *et al*, 2008). However, the complete deafness in VGLUT3 KO mouse conflicts with the variable onset and progression rate of DFNA25. The rare variant VGLUT3-p.A211V was identified as the cause of DFNA25 (Ruel *et al*, 2008). This variation does not impair the glutamate vesicular accumulation, neither the quantal release nor the quantal content in hippocampal autaptic neurons (Ramet *et al*, 2017). In addition, investigation of a mouse line harboring the p.A221V allele in human (p.A224V in mouse, VGLUT3^A224V/A224V^ mouse) showed that this variant resulted to a strong decrease of VGLUT3 expression (≈ −70%) in the brain without altering VGLUT3-piloted behavior (Ramet *et al*, 2017).Thus, it is not so clear whether and how the VGLUT3-p.A211V variant results in the human auditory deficit, DFNA25. To resolve the mechanisms underlying DFNA25, we therefore studied the auditory system of the VGLUT3^A224V/A224V^ mouse line. Our data show that the VGLUT3^A224V/A224V^ mice phenocopied the progressive hearing loss reported in human. The auditory impairment was caused by a selective defect in the IHC stereocilary bundle. In addition, we observed an increase of the ribbon size associated with a larger rate of exocytosis. Taken together, our results indicate unexpected morphological and functional consequences of the VGLUT3-p. A224V variant that underlies DFNA25. Therefore, our VGLUT3^A224V/A224V^ mouse line is a valuable asset to understand and treat DFNA25.

## Materiel and methods

Experiments were carried out in accordance with the animal welfare guidelines 2010/63/EC of the European Communities Council Directive regarding the care and use of animals for experimental procedures. Animals were housed in facilities accredited by the French “Ministère de l’Agriculture et de la Forêt” (Agreement C-34-172-36) and the experimental protocol was approved (Authorization CEEA-LR-12111) by the Animal Ethics Committee of Languedoc-Roussillon (France).

### Animals

We studied A224V mice of either sex, which have been previously described (Ramet *et al*, 2017). In brief, the p.A211V mutation has been isolated in two unrelated human families (Ruel *et al*, 2008). In mouse, the corresponding alanine is at position 224. A mouse line expressing the p.A224V mutation was generated at Phenomin–Institut Clinique de la Souris Illkirch, France; http://www.phenomin.fr/ and was named VGLUT3^A224V/A224V^ as previously reported (Ramet *et al*, 2017). A point mutation was introduced in exon 5 of the mouse Slc17a8 gene: a GCG (coding for an alanine) was exchanged for a GTG (coding for a valine). Mice were bred in-house and maintained on a C57Bl6/J genetic background. Heterozygous mice were bred to generate VGLUT3^A224V/A224V^ and wild-type littermates.

### Genotyping

Genotypes of mice was determine by Polymerase Chain Reaction (PCR) analysis of genomic DNA using FastStart PCR Master Mix (Roche Applied Science, Penzberg, Germany) as previously described (Ramet *et al*, 2017). In brief, tail or toe biopsies were digested overnight at 55°C in 300 µl of lysis buffer (containing in mM: 100 Tris-HCL pH 8.5; 5 EDTA; 0.2% SDS; 200 NaCl; pH : 8.5) with 100µg/ml of proteinase K (Promega, Madison, WI, USA). Samples were centrifuged 5 min at 10 000g and supernatants were collected. DNA was precipitated by addition of 500 µl of isopropanol. Samples were centrifuged 10 min at 10 000 g and supernatants were discarded. DNA was washed with 500 µl of EtOH 70% and centrifuged 5 min at 10 000 g. After evaporation of the EtOH, DNA was suspended in 100 µl of water. The PCR was conducted with the following thermal cycle program: 1 cycle of 95°C for 10 min; 40 cycles of 94°C for 40 s, 62°C for 30 s, 72°C for 1 min; final elongation step at 72°C for 7 min. The PCR primers were the following: p1, 5’-CGGAGGGGAAGCCAGGAAAGGG-3’, and p2, 5’-GACAGCTCAGTGAGCTGTAGACCCAG-3’ for the WT and the mutated allele, yielding bands of 219 and 306 bp, respectively, which were visualized on a 2% agarose gels.

### *In vivo* recordings

Mice were anesthetized by an intraperitoneal injection of a mixture of Zoletil 50 (40 mg/kg) and Rompun 2% (3 mg/kg). The rectal temperature was measured with a thermistor probe and maintained at 37.1°C ± 1°C, using a heated underblanket. (Homeothermic Blanket Systems, Harvard Apparatus). Heart rate was monitored via EKG electrodes.

### Auditory brainstem response and distortion product otoacoustic emission recordings

For auditory brainstem response (ABR), the acoustical stimuli consisted of 10-ms tone bursts, with a 8-ms plateau and 1-ms rise/fall time, delivered at a rate of 20.4/s with alternate polarity by a JBL 2426H loudspeaker in a calibrated free field. Stimuli were presented by varying intensities from 80 dB SPL to 0 dB SPL, in 10 dB step. Stimuli were generated and data acquired using MATLAB (MathWorks, Natick, MA, USA) and LabVIEW (National Instruments, Austin, TX, USA) software. The potential difference between vertex and mastoid intradermal needles was amplified (5000 times, VIP-20 amplifier), sampled (at a rate of 50 kHz), filtered (bandwidth of 0.3-3 kHz) and averaged (600 times). Data were displayed using LabVIEW software and stored on a computer (Dell T7400). ABR thresholds were defined as the lowest sound intensity, which elicits a clearly distinguishable response. For distortion product otoacoustic emission (DPOAE) recordings, an ER-10C S/N 2528 probe (Etymotic Research), consisting of two emitters and one microphone, was inserted in the left external auditory canal. Stimuli were two equilevel (65 dB SPL) primary tones of frequency f1 and f2 with a constant f2/f1 ratio of 1.2. The distortion 2f1-f2 was extracted from the ear canal sound pressure and processed by HearID auditory diagnostic system (Mimosa Acoustic) on a computer (Hewlett Packard). The probe was self-calibrated for the two stimulating tones before each recording. f1 and f2 were presented simultaneously, sweeping f2 from 20 kHz to 2 kHz by quarter octave steps. For each frequency, the distortion product 2f1-f2 and the neighboring noise amplitude levels were measured and expressed as a function of f2.

### Patch-clamp recordings

After cervical dislocation of mice (6 months of age), IHCs of the apical coil of freshly dissected organs of Corti were patch-clamped at their basolateral face at room temperature in tight perforated-patch configurations. The dissection solution contained the following in mM: 5.36 KCl, 141.7 NaCl, 1 MgCl_2_-6H_2_O, 0.5 MgSO_4_-7H_2_O, 10 HEPES and 10 D-glucose. For Ca^2+^ current and capacitance measurement recordings, the extracellular solution contained the following in mM: 2.8 KCl, 105 NaCl, 1 MgCl_2_-6H_2_O, 2 CaCl_2_, 10 HEPES, 35 TEA-Cl, 1 CsCl, 10 D-Glucose. The pipette solution for perforated patch-clamp recordings of Ca^2+^ currents contained the following in mM: 135 Cs-glutamic acid, TEA-Cl, 10 4-AP, 1 MgCl_2_-6H_2_O, 10 HEPES and 400 µg/ml amphotericin B. Solutions were adjusted to pH 7.3 and had osmolarities between 290 and 310 mOsm/kg H_2_O. All chemicals were obtained from Sigma (St. Louis, MO, USA) with the exception of amphotericin B (Calbiochem, La Jolla, USA). Patch pipettes were pulled from borosilicate glass capillaries (Kwik Fil, WPI, Worcester, MA, USA) with a two-step vertical puller PIP 6 (HEKA Elektronik, Lambrecht, Germany) and coated with silicone elastomer (Sylgard).

### Ca^2+^ current recordings

Currents were low-pass filtered at 5 kHz and sampled at 10 kHz for exocytic cell membrane capacitance change (ΔCm) and at 40 kHz for Ca^2+^ current recordings. Ca^2+^ current was isolated using P/n protocols (10 leak pulses with amplitudes of 20% of the original pulse from a holding potential of −117 mV). Cells that displayed a membrane current exceeding −50 pA at −87mV were discarded from the analysis. No Rs compensation was applied, but recordings with Rs > 30 MOhm were discarded from the analysis. Mean resting capacitance were 7.5 ± 0.5 pF and 6.3 ± 0.8 pF for IHCs from VGLUT3^+/+^ and VGLUT3^A224V/A224V^ mice, respectively. All voltages were corrected for liquid junction potentials calculated between pipette and bath (−17 mV).

### Capacitance measurement recordings

Cell membrane capacitance (Cm) was measured using the Lindau–Neher technique (Lindau & Neher, 1988), implemented in the software-lockin module of Patchmaster (HEKA Elektronik) combined with compensation of pipette and resting cell capacitances by the EPC-10 (HEKA Elektronik) compensation circuitries. A 1 kHz, a 70 mV peak-to-peak sinusoid was applied about the holding potential of −87 mV. ΔCm was estimated as the difference of the mean Cm over 400 ms after the end of the depolarization (the initial 250 ms was skipped) and the mean prepulse capacitance (400 ms). Mean ΔCm estimates present grand averages calculated from the mean estimates of individual IHCs.

### Immunohistochemistry, confocal and super-resolution microscopy

Immunohistochemistry was performed on whole-mount preparations of organs of Corti. Mice were decapitated after rapid cervical dislocation and their cochleas were removed from the temporal bone and dissected in the phosphate buffered saline (PBS, containing in mM: 130 NaCl, 2.68 KCl, 10 Na_2_HPO_4_, 1.47 KH_2_PO_4_, pH 7.4) solution. The mid-apical cochlear turns were then fixed 15 min in 4% paraformaldehyde diluted in PBS; afterwards, they were immuno-histochemically processed as a whole-mount. The tissues were rinsed 3×10 min in PBS, then preincubated 1 hour in blocking solution (20% goat serum, 0.3% Triton), then incubated overnight at 4°C in the incubation solution (1% goat serum, 0.1% Triton) with primary antibodies or antisera. The primary antibodies used, and their respective dilutions were CtBP2 1:500 (BD Transduction Laboratories, San Jose, CA, USA; Cat No.:612044), Homer1 1:300 (Merck Millipore, Cat. No. ABN37) and VGLUT3 1:300 (Synaptic Systems, Cat.No. 135 204). Tissues were then rinsed 3×10 min in wash buffer (containing in mM: 15.8 Na2HPO4, 2.9 NaH2PO4, 0.3% Triton X-100, 450 NaCl) and incubated for 2 hours in the incubation solution with fluorescently labeled secondary antibodies, rhodamin-phalloidin (Molecular probes, Eugene, OR, USA) and Hoechst dye (Invitrogen, Carlsbad, CA, USA). They were finally rinsed 4×10 min in wash buffer and mounted in DAKO florescence mounting medium (code-s3023, Agilent Technologies, Inc. Santa Clara, CA, USA). Tissues were examined with the Zeiss LSM880 (Zeiss, Oberkochen, Germany) airyscan or confocal microscopes of the Montpellier RIO Imaging facility (Montpellier, France). Image stacks were then processed with ImageJ software (Wayne Rasband, National Institutes of Health, USA). The quantifications of ribbons and synapses, i.e., juxtaposed spots of the presynaptic ribbon component RIBEYE and postsynaptic density protein Homer, were performed in Z-stacks confocal images using a 3D custom algorithm (Bourien et al, 2014).

For Stimulated Emission Depletion (STED) microscopy imaging, secondary goat anti-guinea pig Alexa-488 (Cat.-Nr. A11039; Thermo Fisher Scientific), goat anti-mouse IgG, Abberior STAR RED (STRED-1001-500UG, Abberior GmbH) and goat anti-rabbit IgG, Abberior STAR-580 (ST580-1002-500UG, Abberior GmbH) antibodies were applied for 2 h at room temperature. After mounting the specimen in Dako mounting medium, image acquisition was performed using an Abberior Instruments Expert Line STED microscope (Abberior Instruments GmbH; based on an Olympus IX83 inverted microscope) in confocal and/or STED mode using a 1.4 NA UPlanSApo 100x oil immersion objective lens. We employed two pulsed lasers 580 nm (red) and 647 nm (far-red) for excitation and a pulsed 775 nm laser for stimulated emission depletion. Image stacks were acquired with Imspector Software, keeping xy pixel sizes of 30 x 30 nm and step sizes of 50 nm (3D-STED). xy pixel sizes in 2D-STED was kept at 30 x 30 nm. Microscope is housed at Montpellier Resource Imaging (MRI) facility (Montpellier, France). Image stacks were then processed using ImageJ (FiJi) software (Wayne Rasband, National Institutes of Health, USA) and ribbon size analysis was achieved by using custom written script in the MATLAB software over 30 IHCs (n=4 cochleas) at 2 months, 73 IHCs (n=4 cochleas) at 4 months, 61 IHCs (n=4 cochleas) at 6 months from VGLUT3^+/+^ mice and 39 IHCs (n=3 cochleas) at 2 months, 118 IHCs (n=3 cochleas) at 4 months, 83 IHCs (n=4 cochleas) at 6 months from VGLUT3^A224V/A224V^ mice. Measurement of ribbon size was obtained by i) surrounding the ribbon using *contour* function (iso-intensity line fixed at mean background + 2 SD) ii) quantifying length of long axis (*a*_lg_) and short axis (*a*_sh_) using the *fit_ellipse* function (least-squares criterion). The elliptic surface (*S*_el_) were then calculated using the equation *S*_el_ =(*a*_lg_×*a*_sh_×π)/4.

### Electron microscopy

Scanning (SEM) and transmission (TEM) electron microscopy were done for morphologic evaluation of the cochlear hair cells. For both techniques, the animals were decapitated under deep anaesthesia (pentobarbital, 50 mg/kg), their cochleas were removed from the temporal bone. For SEM, PBS washed cochlea were fixed with 2.5% glutaraldehyde in phosphate buffer, pH 7.2 for 30 minutes at room temperature, followed by washing in phosphate buffer. The bony capsule of the cochlea was dissected out, and the stria vascularis as well as the tectorial and Reissner’s membranes were removed. Fixed cochleae were dehydrated using a graded ethanol series (30-100%), followed by critical point drying with CO_2_. Subsequently, the samples were sputter coated with an approximatively 10 nm thick gold film and then examined under a scanning electron microscope (Hitachi S4000) using a lens detector with an acceleration voltage of 10kV at calibrated magnifications. For 6-month-old VGLUT3^+/+^ and VGLUT3^A224V/A224V^ mice, 3 cochleas were processed and examined. Examinations were made all along the cochlear spiral (around 6 mm long) from the apex to the basal end. SEM quantification has been carried-out over 1120 IHCs and 3568 OHCs in 6-months VGLUT3^+/+^ (n=4 cochleas) and 956 IHCs and 3226 OHCs 6-months VGLUT3^A224V/A224V^ mice (n=4 cochleas). Mean estimates present grand averages calculated from the mean estimates of individual cochleas.

For TEM, the cochlea were perfused with a solution of 2.5% glutaraldehyde in PHEM buffer (1X, pH 7.4) and immersed in the same fixative for 1 h at room temperature, consequently overnight at 4°C. Sample were then rinced in PHEM buffer and post-fixed in a 0.5% osmic acid for 2 hours at dark and room temperature. After two rinces in PHEM buffer, the cells were dehydrated in a graded series of ethanol solutions (30-100%). The cells were embedded in EmBed 812 using an Automated Microwave Tissue Processor for Electronic Microscopy, Leica EM AMW. Thin sections (70 nm; Leica-Reichert Ultracut E) were collected at different levels of each block. These sections were counterstained with uranyl acetate 1.5% in 70% Ethanol and lead citrate and observed using a Tecnai F20 transmission electron microscope at 200KV. Both SEM and TEM were carried-out at the CoMET facility (MRI, INM, Montpellier, France).

### Data analysis

Data were analyzed using MATLAB and Igor Pro (WaveMetrics, Lake Oswego, OR, USA) softwares. All these data are expressed as means ± standard error of mean (s.e.m.) and were compared by two-tailed Mann-Whitney Wilcoxon test or Student t-test.

## Results

### VGLUT3^A224V/A224V^ mouse mimics DFNA25

We first examined the synchronous activation of the ascending auditory system of the VGLUT3^A224V/A224V^ mice and VGLUT3^+/+^ littermates. In response to incoming sound stimulation, VGLUT3^A224V/A224V^ mice showed a reduction in the auditory brainstem response (ABR) amplitude from 2 months up to 6 months (Mean wave 1 ABR amplitude at 2 months: 1.8 ± 0.27 µV vs 0.6 ± 0.07 µV in VGLUT3^+/+^ and VGLUT3^A224V/A224V^ mice, respectively, p=0.0002; at 6 months: 0.85 ± 0.1 µV vs 0.18 ± 0.05 µV in VGLUT3^+/+^ and VGLUT3^A224V/A224V^ mice, respectively, p=0.0003; **Fig. 1A-D**). The alteration in the ABR waveform was associated with a progressive threshold shift (Mean threshold at 2 months: 31 ± 1.1 dB SPL vs 40.4 ± 2 dB SPL in VGLUT3^+/+^ and VGLUT3^A224V/A224V^ mice, respectively, p= 0.003; at 6 months: 43.5 ± 1.7 dB SPL vs 67.7 ± 1.2 dB SPL in VGLUT3^+/+^ and VGLUT3^A224V/A224V^ mice, respectively, p=1.10^−6^; **Fig. 1E-H**).

**Figure 1:**
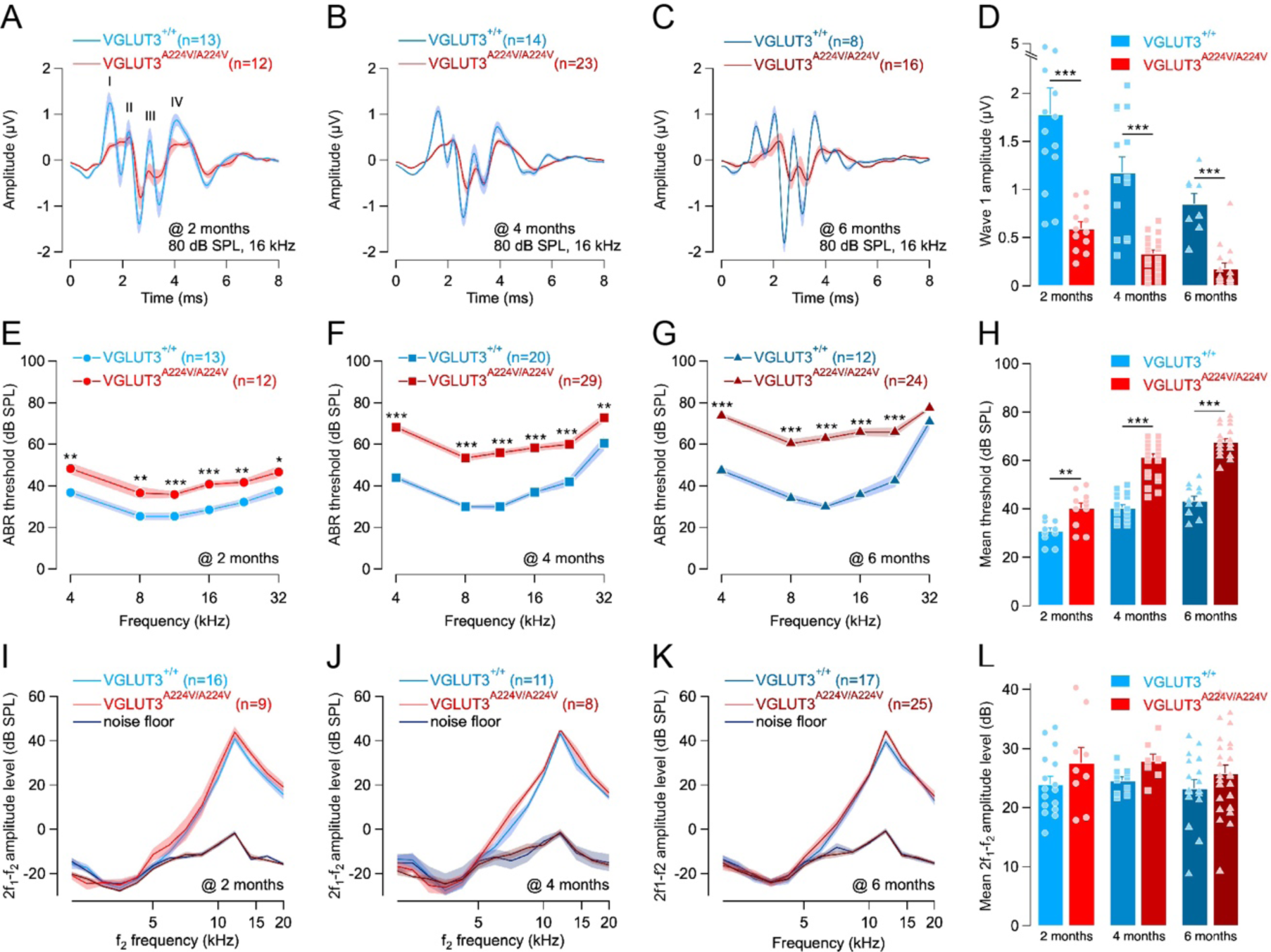
VGLUT3^A224V/A224V^ mouse line mimics human DFNA25 deafness. (**A, C**) Mean ABR recordings evoked by 16-kHz tone burst at 80 dB SPL from 2- to 6-month-old VGLUT3^+/+^ and VGLUT3^A224V/A224V^mice. (**D**) Mean wave 1 ABR amplitude from (**A**-**C**). (**E-G)** Mean ABR audiograms from 2- to 6-month-old VGLUT3^+/+^ and VGLUT3^A224V/A224V^mice. (**H**) Mean threshold from (**E**-**G**). (**I**-**K**) DPOAEs from 2- to 6-month-old VGLUT3^+/+^ and VGLUT3^A224V/A224V^mice. The 2f1-f2 amplitude level is shown as a function of f2 frequency. The black lines indicate the background noise level. (**L**) Mean 2f1-f2 amplitude level from (**I**-**K**) measured between 5 to 20 KHz. n indicates the number of cochleae recorded. Level of significance: *p < 0.05; **p < 0.01; ***p < 0.001, two-tailed Mann-Whitney Wilcoxon test.

To assess the OHCs function, we then probed the distortion product otoacoustic emissions (DPOAEs). We observed robust DPOAEs amplitude in VGLUT3^A224V/A224V^ mice, indicating that OHC activity was essentially preserved (2f_2_-f_1_ @ f_2_ 11.9 kHz: 44.4 ± 1.5 dB SPL vs 39.6 ± 1.8 dB SPL in 6 month-old VGLUT3^+/+^ and VGLUT3^A224V/A224V^, respectively; **Fig. 1I-L**). Altogether, the VGLUT3^A224V/A224V^ mouse replicates progressive hearing loss of DFNA25 with functional cochlear amplification.

### VGLUT3 is normally expressed in VGLUT3^A224V/A224V^ mouse

The variant strongly decreases (−70%) the expression of VGLUT3-p.A224V in the brain (Ramet *et al*, 2017). To determine whether a defect of the VGLUT3 expression might contribute to the threshold shift, we examined the distribution of VGLUT3 in the IHCs from VGLUT3^A224V/A224V^ mouse. In both wild-type and mutant mouse, VGLUT3 was distributed throughout the cell, from the upper side of the nucleus to the basolateral side (**Fig. 2 A, B)**. Semi-quantitative analysis of the immunofluorescence showed similar distribution and expression of VGLUT3 between both genotypes (VGLUT3 immunofluorescence: 1710 ± 26 A.U. vs 1678 ± 49 A.U.xµm in IHCs from 4-month-old VGLUT3^+/+^ and VGLUT3^A224V/A224V^ mice, respectively; **Fig. 2 C**). Therefore, in contrast to what is observed in the central nervous system neurons (Ramet *et al*, 2017), the p.A22AV variant does not disturb the VGLUT3 expression and targeting within the hair cells cytoplasm.

**Figure 2:**
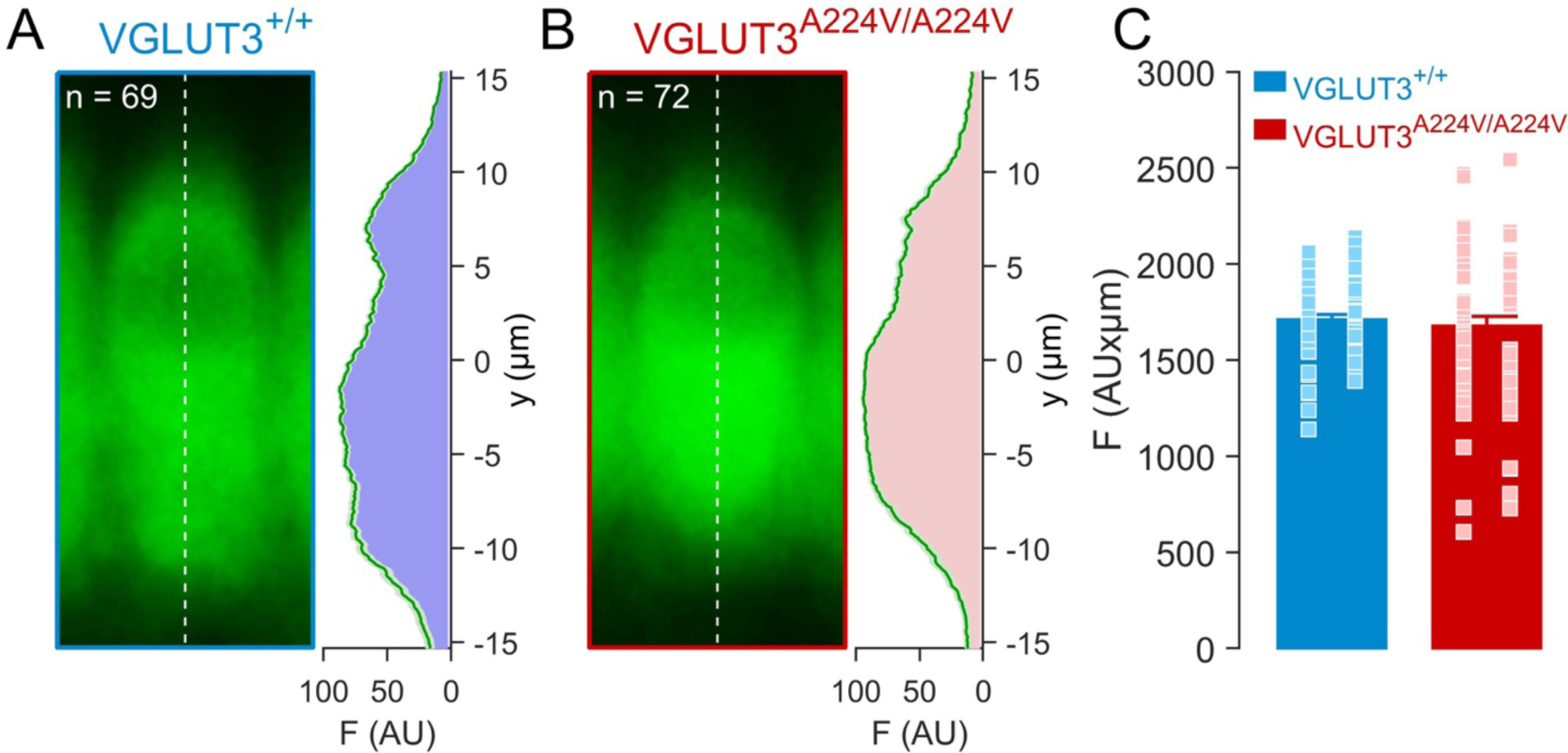
VGLUT3 distribution in IHCs from VGLUT3^+/+^ and VGLUT3^A224V/A224V^ mice. (**A, B**) Maximum projections average of confocal sections of 4-month-old VGLUT3^+/+^ (**A**) and VGLUT3^A224V/A224V^ (**B**) mid-apical turns IHCs immunolabelled against VGLUT3 (green) processed and imaged under identical conditions. Left panel: IHCs have been centered in respect to the centroid of the nucleus; right panel: average of the maximum projections fluorescence measured through the vertical central axis crossing the IHCs (dashed white line). n indicates the number of IHCs immunolabelled (n= 4 cochleas for each genotype). (**C**) Semiquantitative analysis of VGLUT3 immunofluorescence. Bar histograms represent the integral of the maximum projection fluorescence measured through the vertical central axis crossing the IHCs of 4-month-old VGLUT3^+/+^ (blue) and VGLUT3^A224V/A224V^ (red) corresponding to (**A, B**), respectively.

### Alteration at the stereociliary bundle in VGLUT3^A224V/A224V^ mouse

Alteration of the ABR together with intact DPOAEs is the hallmark of auditory neuropathy. We therefore examined the morphology of the inner hair cells. Surprisingly, we found-out disarrayed or fused stereocilia, in 3-month-old VGLUT3^A224V/A224V^ mice contrasting with the clearly individual and staircase pattern of the stereocilia bundle in the VGLUT3^+/+^ mouse (**Supplementary data 1**). To examine the defect of the IHCs at a higher resolution, we used scanning electron microscopy. While apical pole of the IHCs in VGLUT3^+/+^ mice shows a normal appearance, with staircase-organized stereocilia, we observed fused stereocilia bundles in the IHCs from VGLUT3^A224V/A224V^ mouse as early as 1 month of age (**Supplementary data 2**). In 6 month-old VGLUT3^A224V/A224V^ mice, disorganized or even absent cilia apparatus were found in a large fraction of IHCs at the apical region of the cochlea, e.g., the frequency-encoding region up to 20 kHz (apical turn: 17.4 ± 4.1 % and 58.2 ± 11 % of IHC with altered stereocilia bundle in VGLUT3^+/+^ and VGLUT3^A224V/A224V^ mice, respectively, p=0.03, **Fig. 3 A, B**). In contrast, the fraction of IHC harboring fused stereociliary bundle drastically drops at the basal end of the cochlea, corresponding to the 32-50 kHz frequency-encoding region (basal turn 4.8 ± 2.1 % and 18.8 ± 6.3 % of IHC with altered stereocilia bundle in VGLUT3^+/+^ and VGLUT3^A224V/A224V^ mice, respectively, **Fig 3 C, D**). Consistent with preserved DPOAEs, the stereocilia looked pretty normal in the OHCs along the entire tonotopic gradient (apical turn: 92 ± 1.7 % and 91.2 ± 2.7 % of OHC with normal stereocilia bundle in VGLUT3^+/+^ and VGLUT3^A224V/A224V^ mice, respectively, **Fig. 3 A, B**; basal turn: 96.7 ± 0.1 % and 91.8 ± 5.4 % of OHC with normal stereocilia bundle in VGLUT3^+/+^ and VGLUT3^A224V/A224V^ mice, respectively; **Fig. 3 C, D**). Together, these data suggest that the threshold shift in VGLUT3^A224V/A224V^ mouse stems from defective stereocilia architecture of the IHCs.

**Figure 3:**
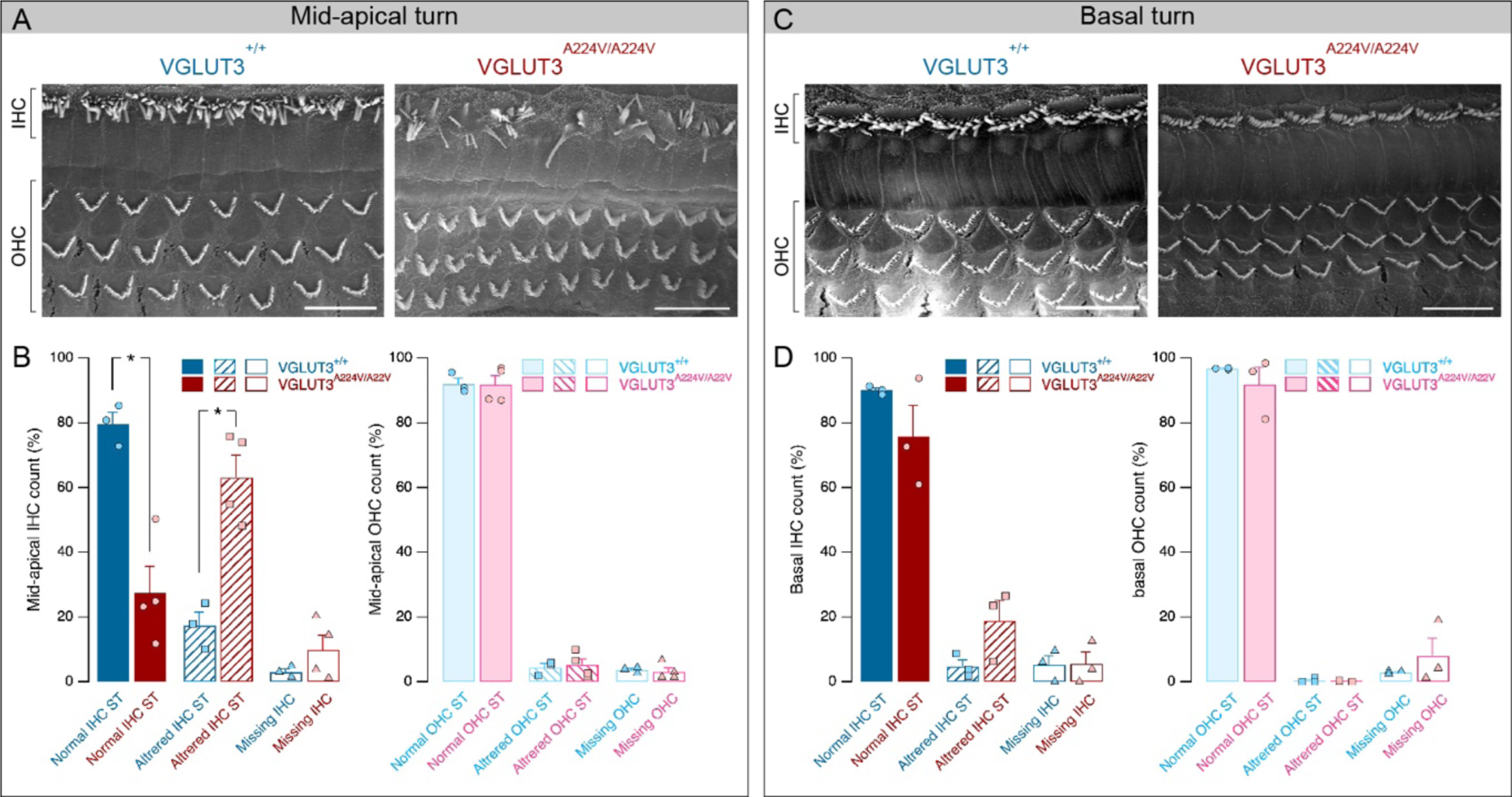
Stereociliary bundle morphology in hair cells at 6 months. (**A, C**) SEM of the organ of Corti from mid-apical (**A**) and basal (**C**) turns in 6-month-old VGLUT3^+/+^ and VGLUT3^A224V/A224V^ mice. (**B**) Fraction of the IHCs (left) and OHCs (right) harboring normal or altered stereocilia (ST) and missing hair cells observed at the apical to mid turn (**B**) or basal turn (**D**) of the cochlea. Level of significance: *p < 0.05, two-tailed Mann-Whitney Wilcoxon test. Scale bars: 10 µm.

### Long-term alteration in the ribbon synapse morphology

Given the role of VGLUT3 in the synaptic transfer between the IHC and the afferent fiber terminal, we thought that additional consequences may occur toward the pre-synaptic compartment, i.e., the synaptic ribbon. Akin to wild-type mice, the IHCs from 2 months old VGLUT3^A224V/A224V^ mice harbor plasma-membrane anchored synaptic ribbons, surrounded by a halo of vesicles and facing the post-synaptic density of the afferent fiber (**Supplementary data 3 A, C**). At two-months of age, the numbers of immunolabeled ribbons with anti-Ctbp2 antibody were comparable between wild-type and VGLUT3^A224V/A224V^ mice (13.2 ± 1.1 ribbon per IHC vs 13.8 ± 0.6 ribbon per IHC in VGLUT3^+/+^ and VGLUT3^A224V/A224V^ mice, respectively, **Fig 4 A-C**).

**Figure 4:**
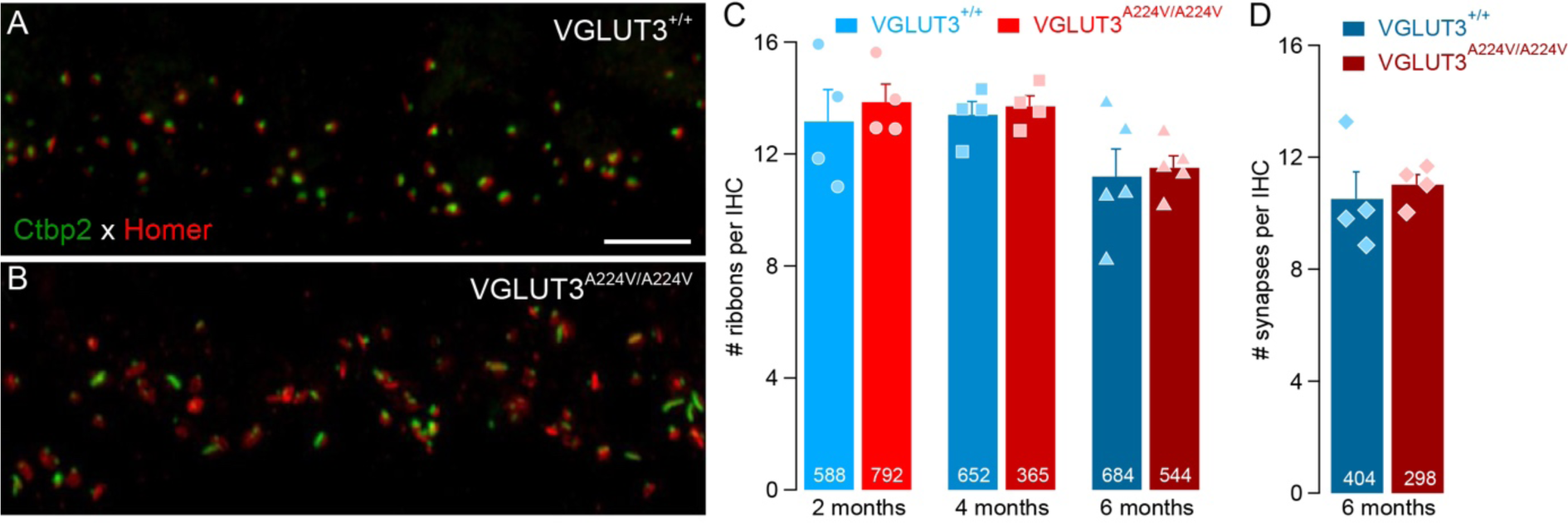
Ribbon and synapses count in IHCs. **(A-B)** Synaptic ribbons and post-synaptic densities immunostained with antibodies against Ctbp2 (green) and Homer (red), respectively, in IHCs from 6 months VGLUT3^+/+^ (**A**) and VGLUT3^A224V/A224V^ (**B**) mice. Scale bar: 5 µm. (**C**) Mean of ribbons per IHC given by the immunostaining of Ctbp2 from 2-, 4- and 6-months VGLUT3^+/+^ and VGLUT3^A224V/A224V^ mice. (**D**) Mean of synapses per IHC given by the juxtaposed immunostaining of Ctbp2 and Homer from 6 months IHCs VGLUT3^+/+^ and VGLUT3^A224V/A224V^ mice. White numbers indicate the number of IHCs analyzed. For each age and genotype, 4 cochleas have been used except for 6 months counting in **C (**n=5 cochleas).

To obtain a quantitative size analysis over a large number of ribbons, we observed the synaptic bodies using STED superresolution microscopy (**Fig. 5**). Ctbp2-immunolabeled signals looked like ovoid spots with almost undistinguishable size between both genotypes (long object axis: 440.3 ± 11.3 nm vs 458.2 ± 8.7 in VGLUT3^+/+^ and VGLUT3^A224V/A224V^ mice, respectively; short object axis: 287.4 ± 4.3 in VGLUT3^+/+^ and 302.4 ± 3.8 VGLUT3^A224V/A224V^ mice, respectively, **Fig. 5 A-C**). Because the VGLUT3^A224V/A224V^ mouse already shows a threshold shift at 2 months of age, these results exclude any morphological changes at the presynaptic compartment as the primary cause for the VGLUT3^A224V/A224V^ mouse hearing loss.

**Figure 5:**
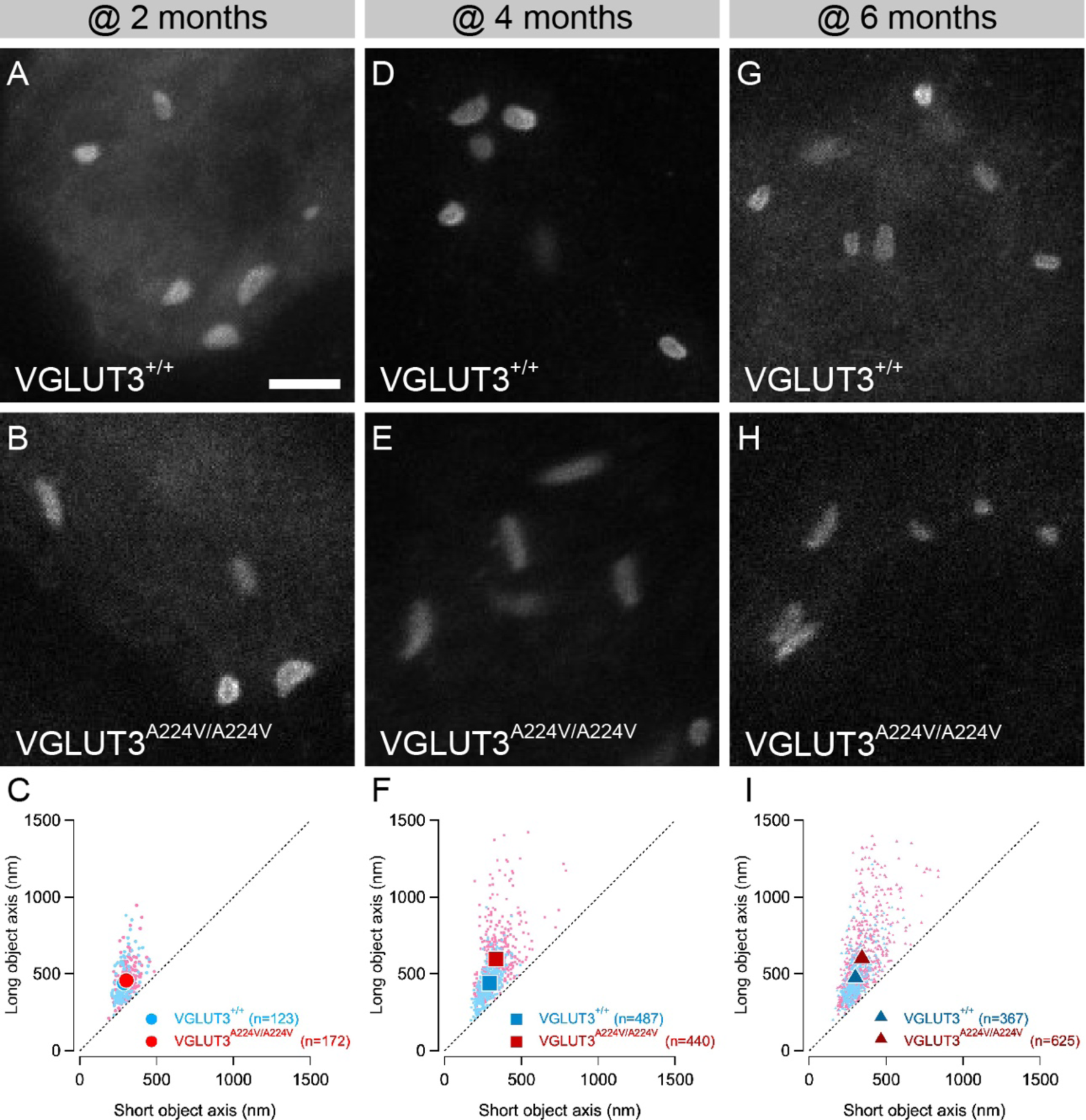
Ribbon synapse morphology in the IHCs. **(A-B, D-E** and **G-H)** Representative STED images of fluorescently labeled synaptic ribbons of 2 months (**A, B**), 4 months (**D, E**) and 6 months (**G, H**) IHCs from VGLUT3^+/+^ (**A, D** and **G**) and VGLUT3^A224V/A224V^ (**B, E** and **H**) mice. Scale bar: 1 µm. (**C, F** and **I**) Long versus short axes for VGLUT3^+/+^ (blue) and VGLUT3^A224V/A224V^ (red) ribbons at 2-months (**C**), 4-months (**F**) and 6-months (**I**). n indicates the number of ribbon examined.

At 4 and 6 months of age, we still observed ribbon surrounded by halo of synaptic vesicle using transmission electron microcopy (**Supplementary data 3 B, D**). In addition, we did not notice any difference in the ribbon synapse numbers corresponding to the juxtaposed immunostainings of Ctbp2 and post-synaptic scaffolding protein Homer (@ 6 months: 10.5 ± 0.9 ribbon per IHC vs 11 ± 0.7 ribbon per IHC in VGLUT3^+/+^ and VGLUT3^A224V/A224V^ mice, respectively, **Fig 4 D**). However, synaptic ribbons clearly became distorted in the knock-in mice (**Fig 4 A, B**). Using super-resolution microscopy, we could quantify a drastic elongation of the ribbons in the IHCs from VGLUT3^A224V/A224V^ mice at 4- and 6-months-old (@ 6 months: long object axis: 473.7 ± 7.4 nm vs 602.2 ± 11.4 in VGLUT3^+/+^ and VGLUT3^A224V/A224V^ mice, respectively, p<0.001; short object axis: 300 ± 3.1 in VGLUT3^+/+^ and 344.4 ± 4.4 VGLUT3^A224V/A224V^ mice, respectively, p<0.001; **Fig. 5 D-I**).

Consistently, mean elliptic area calculated from ribbon long and short axes shows that the size of the ribbons is similar at 2 months of age in both genotype (Elliptic size: 0.102 ± 0.004 µm^2^ vs 0.111 ± 0.003 µm^2^ in 2-month-old VGLUT3^+/+^ and VGLUT3^A224V/A224V^ mice, respectively, **Fig. 6 A**) but increases at 4 and 6 months in the IHCs form VGLUT3^A224V/A224V^ mice (Elliptic size: 0.116 ± 0.003 µm^2^ vs 0.18 ± 0.006 µm^2^ in 6-month-old VGLUT3^+/+^ and VGLUT3^A224V/A224V^ mice, respectively, p<0.001, **Fig. 6 A**). Size distributions were skewed for 4- and 6-months VGLUT3^A224V/A224V^ mice (**Fig 6 B**). Cumulative size histogram showed that IHCs from 6 months VGLUT3^A224V/A224V^ mice contain ribbons resembling those found at 2 months as well as ribbons for which size increases up to 100 % (Ribbon size up to 0.064 µm^2^ and 0.065 µm^2^ within the 10^th^ percentile against 0.17 µm^2^ and 0.35 µm^2^ within the 90^th^ percentile from 6 months IHCs VGLUT3^+/+^ and VGLUT3^A224V/A224V^ mice, respectively, **Fig. 6 C**). These results indicate that a population of pre-synaptic ribbon undergoes a structural change in the VGLUT3^A224V/A224V^ mouse line.

**Figure 6:**
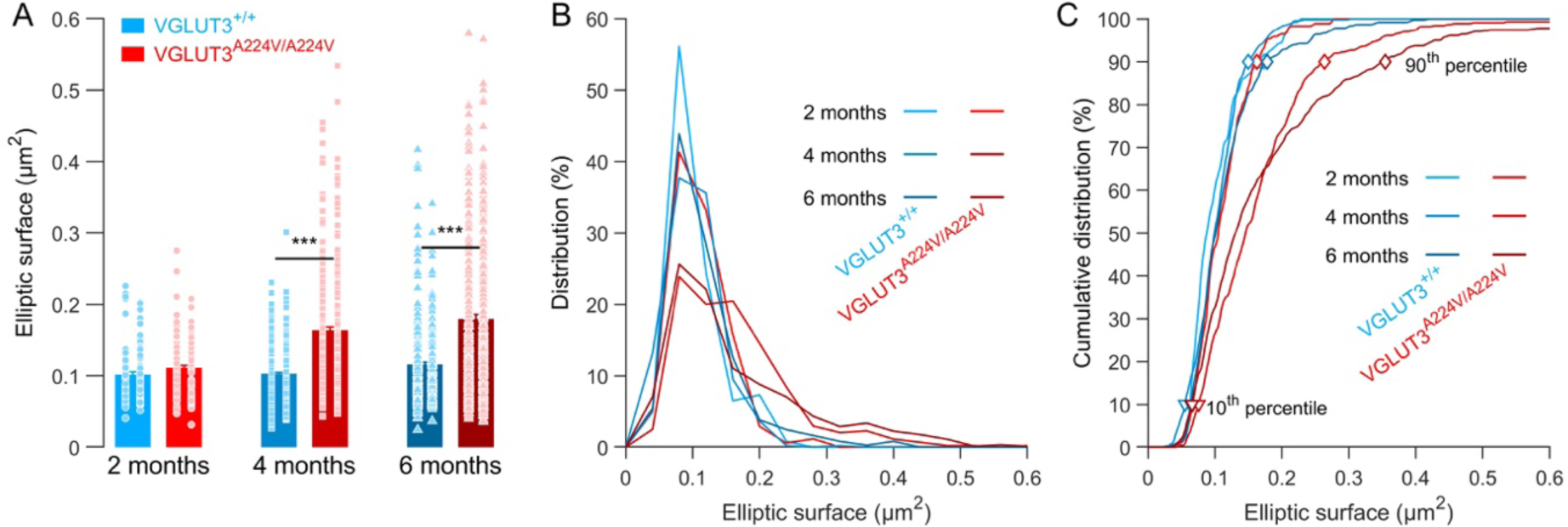
Increase of the elliptic surface of ribbon’s section in the IHCs. **(A)** Mean elliptic surface calculated from the ribbon long and short axes length of 2, 4 and 6 months IHCs from VGLUT3^+/+^ and VGLUT3^A224V/A224V^. (**B**) Distribution of the ribbons plots against the elliptic surface estimates of their cross-section derived from STED-imaging (bin width = 0.04 µm^2^). (**C**) Cumulative distribution of the ribbons as a function of the elliptic surface (bin width = 0.0001 µm^2^). Open symbols indicate the 10^th^ (triangle) and 90^th^ (diamond) percentile. Level of significance: ***p < 0.001, Student test for independent samples.

### Calcium-triggered exocytosis in 6 month-old IHCs VGLUT3^A224V/A224V^ mouse

To determine whether changes in the morphology would affect synaptic release, we probed the calcium-triggered exocytosis at 6 months using patch-clamp membrane capacitance measurements (**Fig. 7 A, B**). The calcium current-voltage relationship did not substantially differ between the genotypes (calcium peak current I_Ca_: −131.5 ± 4.9 pA vs −145.4 ± 15.6 pA for VGLUT3^+/+^ and VGLUT3^A224V/A224V^, respectively, **Fig. 7 C**). Although the exocytosis in IHCs evoked by short time duration depolarizing voltage step (under 20 ms) remains identical (ΔCm_20ms_ 8.4 ± 1.6 fF vs 11.9 ± 2.2 fF for VGLUT3^+/+^and VGLUT3^A224V/A224V^, respectively), we observed a larger increase in the rate of exocytosis from VGLUT3^A224V/A224V^ mice in response to long depolarizing voltage steps (between 50 up to 200 ms time duration; ΔCm_50-_ 200ms 624.7 ± 69.8 fF.s^-1^ vs 320.2 ± 77 fF.s^-1^ for VGLUT3^+/+^ and VGLUT3^A224V/A224V^, respectively; p= 0.0043) (**Fig. 7 D**). Thus, the p.A224V variant of the *SLC17A8* gene is associated with a change in the presynaptic structure, altering the exocytosis capability in hair cells.

**Figure 7:**
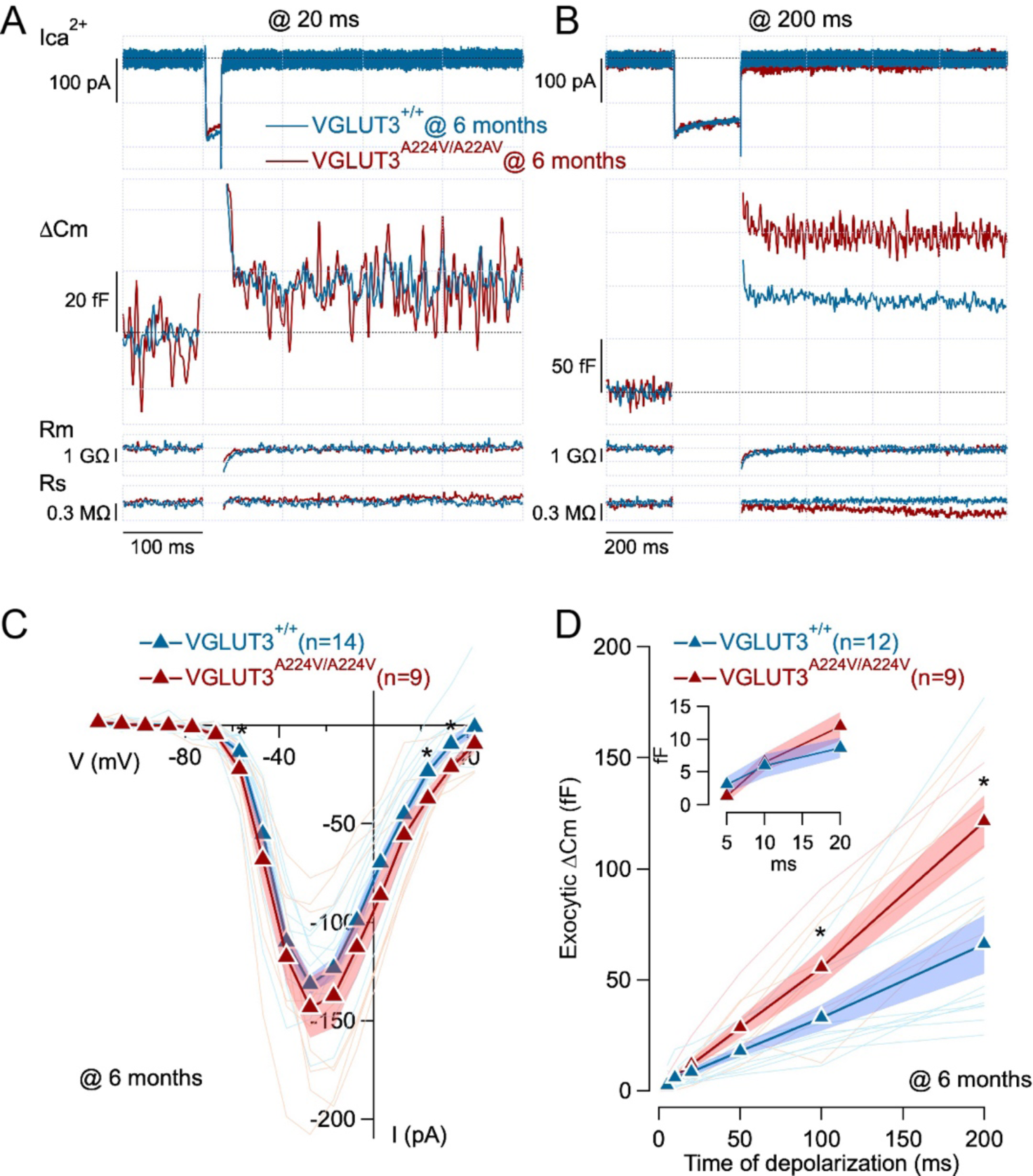
Increase of the sustained releasable pool exocytosis in 6 month-old IHCs VGLUT3^A224V/A224V^ mouse. (**A, B**) Perforated patch-clamp recordings of calcium-triggered exocytosis in 6 month-old IHCs VGLUT3^+/+^ (blue) and VGLUT3^A224V/A224V^ mouse (red). Inward calcium current (ICa^2+^), membrane capacitance jump (ΔCm), membrane resistance (Rm) and series resistance (Rs) evoked by 20 ms (**A**) and 200 ms (**B**) long depolarizing voltage steps from holding potential of −87 mV to −27 mV. (**C**) Ca^2+^current I/V relationships of IHCs from VGLUT3^+/+^ and VGLUT3^A224V/A224V^ mouse. *p < 0.05, two-tailed Mann-Whitney Wilcoxon test. (**D**) Exocytic ΔCm plotted against the duration of depolarization for IHCs from VGLUT3^+/+^ and VGLUT3^A224V/A224V^ mouse. Inset show the membrane capacitance increases in response to depolarizing voltage steps from 5 to 20 ms time duration. Calcium-triggered exocytosis were evoked by depolarizations to −27 mV from holding potential of −87 mV. n: number of IHCs. *p < 0.05, two-tailed Mann-Whitney Wilcoxon test.

## Discussion

In this study, we showed that the mutant mice carrying the p.A224V variant of the *SLC17A8* gene develops a progressive hearing loss, reminiscent of auditory neuropathy. The mutation appears to primarily alter the stereociliary bundle structure of the IHCs followed by a structure-function change in the synaptic ribbon.

### The VGLUT3^A224V/A224V^ mouse as a model of DFNA25

Only few mutations that cause progressive and early-onset form of human deafness DFNA25 have been described (Ryu *et al*, 2016; Ruel *et al*, 2008). Among them, the c.616dupA variant of *SLC17A8* introduces a stop codon and hence leads to a truncated protein (Ryu *et al*, 2016). Another variant is the pA211V allele of *SLC17A8* (Ruel *et al*, 2008). In a virtual 3D model, replacing alanine in position 211 (224 in mouse) by a valine does not substantially modify the VGLUT3 structure as alanine and valine amino acid are closely related (Ramet *et al*, 2017). Here, the mouse line carrying the VGLUT3-p.A224V variant corresponding to the human p.A211V allele shows increasing threshold shift with age. Thus, this result further validates the p.A211V as one of the mutation responsible of DFNA25 (Ruel *et al*, 2008). While otoacoustic emissions failed to be measured in the patients harbouring the p.A211V allele, decent DPOAEs were recorded in the knock-in mutant mouse. This is consistent with the lack of VGLUT3 expression in OHCs and robust DPOAEs measured in the VGLUT3-null mice (Ruel *et al*, 2008; Seal *et al*, 2008). Thus, DFNA25 belongs to the class of the auditory deficit called auditory neuropathy (Moser & Starr, 2016).

### Collapse of the stereocilia bundle in VGLUT3^A224V/A224V^

Up to now, mechanisms underlying DFNA25 were completely unknown. Given that the VGLUT3 KO mouse leads to congenital deafness and that DFNA25 is a progressive deafness (Ruel *et al*, 2008; Seal *et al*, 2008), the p.A211V variant should not lead to a complete loss-of-function behavior of VGLUT3 (Ruel *et al*, 2008). This mutation reduces the amount of VGLUT3 per synapses in the central nervous system neurons (Ramet *et al*, 2017). Therefore, the p.A224V variant may interfere with the expression levels and distribution of VGLUT3 within the hair cells. However, the p.A224V mutation does not disturb the targeting of VGLUT3 in cochlear hair cells. Different transport mechanism of VGLUT3-positive synaptic vesicles may account for the discrepancy between central nervous system neurons and hair cells.

A major alteration lies in the stereocilia architecture. This is somehow surprising since VGLUT3 is absent from IHC stereocilia. In addition, the loss of VGLUT3 does not affect the transducer activity, shown by intact hair cell receptor potential in the absence of VGLUT3 (Obholzer *et al*, 2008; Ruel *et al*, 2008). It has been demonstrated that the membrane trafficking is taking place throughout the hair cell (Kamin *et al*, 2014). Mutated VGLUT3 may lead to a traffic jam of synaptic vesicles at the upper side of the nucleus and destabilize the route for protein turnover toward the stereocilia machinery. This, however, is unlikely as we did not observe any accumulation of VGLUT3 above the nucleus. In addition, the vesicles are most likely recycling at the basal side of the IHCs (Kamin *et al*, 2014; Neef *et al*, 2014; Grabner & Moser, 2018; Kroll *et al*, 2019). The alanine to valine mutation occurs within a cytoplasmic loop that faces the pore of the transporter (Ramet *et al*, 2017). Although the mutation does not seem to affect the structure or primary function of VGLUT3, it might change binding properties with interacting partner. Given our present results, one can hypothesize that VGLUT3 interacts with a partner expressed in IHC and that this protein/protein interaction is also involved into the maintenance of the stereocilia and/or the cuticular plate. Because stereocilia are anchored into the cuticular plate, any defect in its composition would make the stereocilia bundle collapse (Surel *et al*, 2016). Identification of this partner could help to clarify how the VGLUT3 mutation affects stereocilia architecture. This will require furthers investigations, such as isolation of partners interacting with wild-type and p.A211V mutated form of VGLUT3. A detailed examination along the tonotopic axis of the cochlea revealed that the fused stereocilia mostly take place at the apical turn of the cochlea, which are the corresponding region probed in our ABR. In contrast, hair cells from the basal turn seem to be less or not affected by the mutation. This difference might reflect a differential expression of VGLUT3 across the tonotopic axis (Yoshimura *et al*, 2014), the absence of a partner protein in the cochlear basal turn or an activity-dependent alteration. Here again, further experiments are required to dwell on this discrepancy between cochlear turns.

### Synaptic plasticity in VGLUT3^A224V/A224V^

Given that ribbon synapse number and anatomy seemed unaltered in young VGLUT3^A224V/A224V^ (2 months old), ABR reduction and threshold shift are more likely to stem from the defect in the IHC mechano-transduction apparatus. Still, we observe that ribbon synapse changed in their anatomy at later stages. It has been shown that the synaptic ribbon size varies with the calcium influx amplitude (Martinez-Dunst *et al*, 1997; Schnee *et al*, 2005; Frank *et al*, 2009; Regus-Leidig *et al*, 2010; Ohn *et al*, 2016; Sheets *et al*, 2017). However, the lack of change in the calcium current amplitude in IHC of VGLUT3^A224V/A224V^ mice exclude such mechanism. The abnormal shape of ribbons at 4- and 6-months is reminiscent to the thin shape of the ribbons in the absence of VGLUT3 (Seal *et al*, 2008). Thus, these results suggest that the integrity of VGLUT3 is essential for the structure maintenance of the presynaptic ribbon. Here again, the VGLUT3-p.A224V mutation may impede the transport of protein required to adequately shape the ribbon. Interestingly, the elongated ribbons still tether monolayer of synaptic vesicles, indicating that the synaptic vesicle trafficking is VGLUT3-independent.

The IHCs calcium-triggered exocytosis probed by short time duration corresponds to the readily releasable pool (RRP) of synaptic vesicle. In our experiments, the RRP is not affected in the IHCs from VGLUT3^A224V/A224V^ mice. In hair cells, the calcium channels are densely packed underneath the presynaptic element and the RRP correlates to the membrane-proximal synaptic vesicles, i.e., the vesicles tether to the synaptic body and docked to the plasma membrane (Khimich *et al*, 2005; Schnee *et al*, 2005; Rutherford & Roberts, 2006; Meyer *et al*, 2009; Frank *et al*, 2010; Pangršič *et al*, 2010; Wong *et al*, 2014; Neef *et al*, 2018). Thus, the calcium current as well as the RRP size are likely to scale with the number of synapses (Meyer *et al*, 2009). The comparable calcium current amplitude and similar amount of RRP between both genotypes are quite consistent with the preserved number of synapses. The secretion evoked by longer time duration corresponds to the sustained releasable pool (SRP) of synaptic vesicle, which reflects the re-supply of synaptic vesicle towards the release sites (Moser & Beutner, 2000; Pangršič *et al*, 2010; Schnee *et al*, 2011, see for review Moser *et al*, 2019). The secretion associated with the SRP is dramatically increased in mutant mice. This change may be due to a larger number of synaptic vesicles populating the synaptic ribbons as they become elongated in the mutant mouse. In this hypothesis, the synaptic vesicles attached to the ribbon and facing the cytoplasm, the “ribbon-associated SV” may be the morphological correlate to the SRP (Hull *et al*, 2006; Frank *et al*, 2010; Snellman *et al*, 2011; Maxeiner *et al*, 2016). However, this hold true only if synaptic ribbons are still orientated in a perpendicular direction in respect to the plasma membrane. Therefore, extensive 3D reconstruction are required to support this scenario.

What could be the consequence of having an increase in the SRP? It has been proposed that the re-supply of synaptic vesicles build-up the standing RRP *in vivo* and therefore is essential for the neuronal spike rate (Pangršič *et al*, 2010). The increase in the SRP may drive outnumbers of synaptic vesicles and create a traffic jam at the release sites. This is however unlikely as having more available vesicles does not impede the spike rate in the auditory fibers (Pangršič *et al*, 2010). It has been shown that the specificity of the auditory nerve fibers is associated with the pre-synaptic ribbon anatomy. While high-spontaneous rate fibers with low-threshold activation project onto small synaptic ribbons on the pillar side, low-spontaneous rate (LSR) fibers with high-threshold activation connect to large ribbon on the modiolar side (Merchan-Perez & Liberman, 1996; Liberman *et al*, 2011; Ohn *et al*, 2016). Therefore, the increase in the number of thin and elongated synaptic ribbon may reflect an increase in the LSR fibers pool connecting the IHCs in the VGLUT3^A224V/A224V^ mice. In this case, the hearing loss at late stages in VGLUT3^A224V/A224V^ mice would not only be attributable to the stereociliary bundle defect but also to the delayed and high jitter of the first-spike latency within the auditory nerve fibers (Bourien *et al*, 2014).

In conclusion, we showed that the pA211V mutation of *SLC17A8*, the gene encoding VGLUT3, results in a progressive hearing loss resembling to an auditory neuropathy. The p.A224V mutation of VGLUT3 alters both transducer apparatus in the IHCs as well as the synaptic ribbon architecture and release capability. These results highlight the critical role of VGLUT3 in maintaining the hair cell integrity to operate sound-coding. The VGLUT3^A224V/A224V^ mouse will be a valuable asset to fully elucidate molecular mechanisms underlying DFNA25 and to identify a treatment.

## Acknowledgments and Funding sources

The authors thank Chantal Cazevieille for expertise and assistance in electron microscopy. This work was supported by an EU Horizon 2020 Marie Sklodowska-Curie Action Innovative Training network, H2020-MSCA-ITN-2016 [LISTEN - 722098]. SM received a PhD fellowship from the “Ecole de l’INSERM-Liliane Bettencourt”. G.S. is a recipient of the postdoctoral fellowship of University of Montpellier. Authors report no conflict of interest.

## Supplementary information

**Supplementary data 1:**
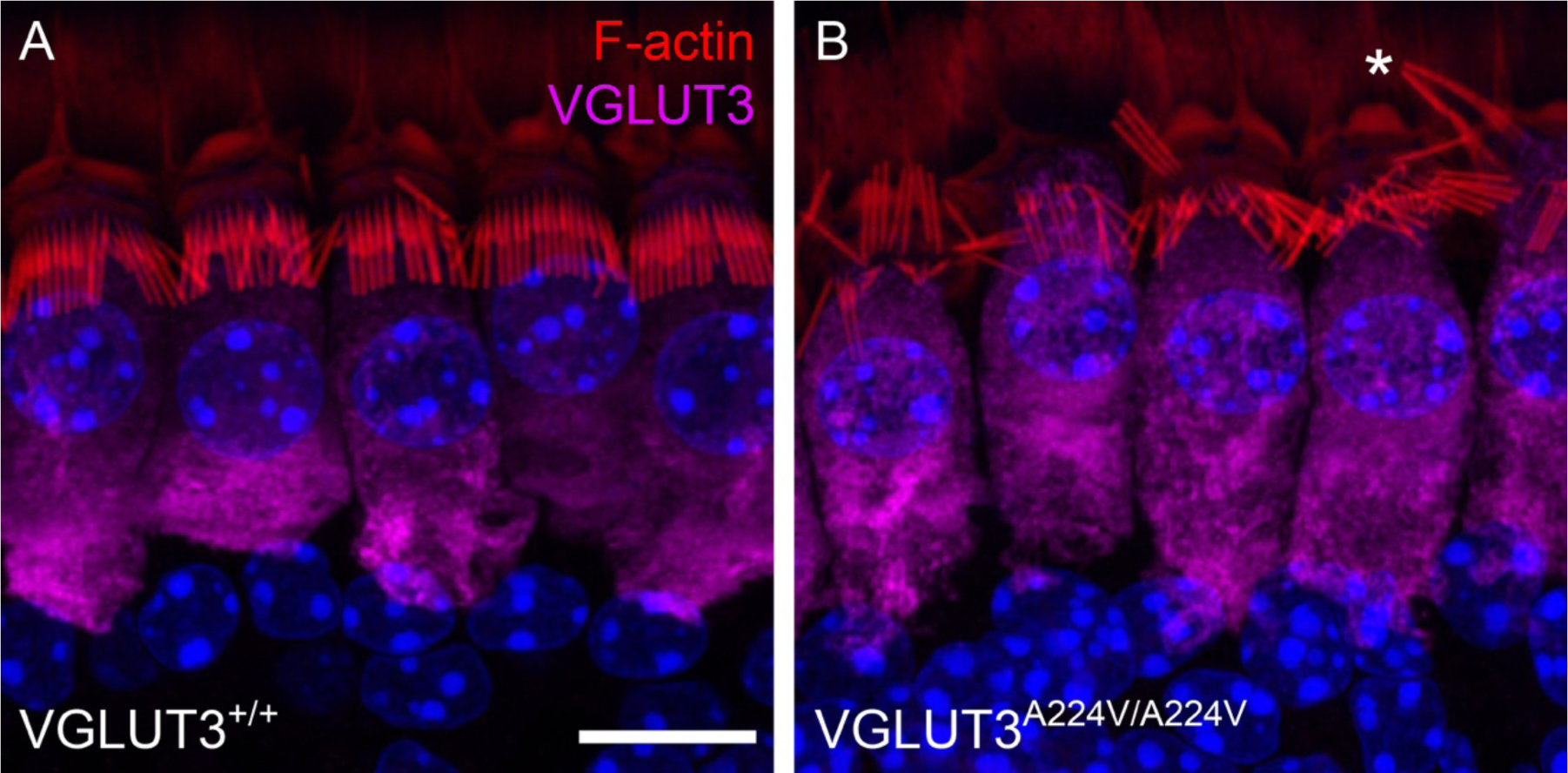
Alteration in the stereocilia bundle of the VGLUT3^A224V/A224V^ mice. (**A**-**B**), F-actin and VGLUT3 distribution at the IHC apical side from 3-month-old VGLUT3^+/+^ (**A**) and VGLUT3^A224V/A224V^ (**B**) mice. Actin filaments are labeled by phalloidin-rhodamine (red) and VGLUT3 is stained using antibody anti-VGLUT3 (purple). Asterisk indicates abnormal fused stereocilia in the VGLUT3^A224V/A224V^ mice. Scale bar: 10 µm.

**Supplementary data 2:**
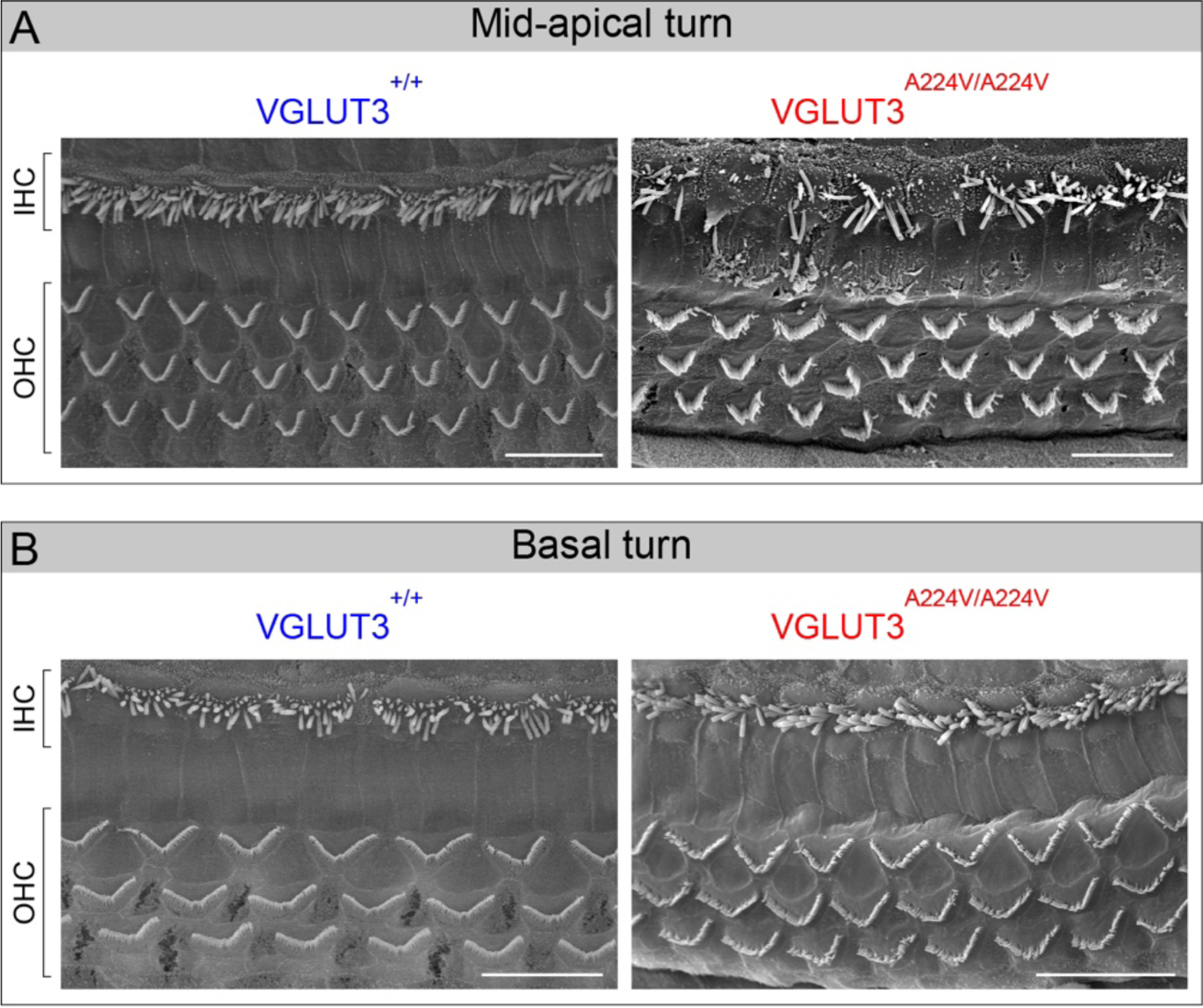
Stereociliary bundle morphology in hair cells at 1 month. (**A, B**) SEM of the organ of Corti from mid-apical (**A**) and basal (**B**) turns in 1-month-old VGLUT3^+/+^ (blue) and VGLUT3^A224V/A224V^ (red) mice. Scale bars: 10 µm.

**Supplementary data 3:**
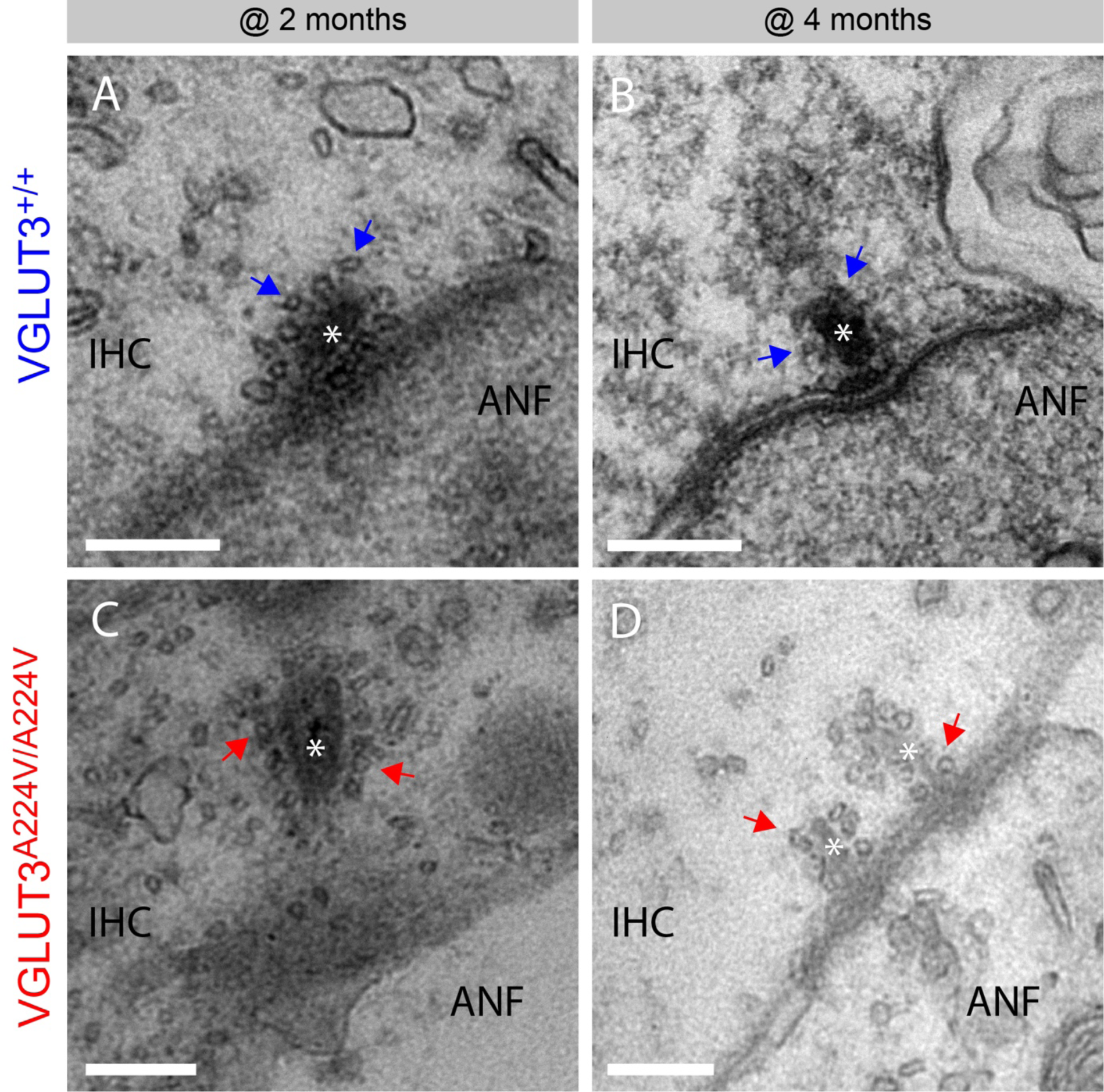
Transmission electron microscopy of synaptic ribbons in IHCs. (**A**-**D**) show examples of synaptic ribbons (white stars) surrounded by a halo of synaptic vesicles (arrows) in IHCs from VGLUT3^+/+^ (**A, B**) and VGLUT3^A224V/A224V^ (**C, D**) at 2 months (**A, C**) and 4 months (**B, D**). ANF: auditory nerve fiber. scale bar: 200 nm.

